# Modeling physiological insulin sensitivity and pre-diabetic insulin resistance in multiple hPSC-derived cell types

**DOI:** 10.1101/2025.10.14.682181

**Authors:** Max Friesen, Rudolf Jaenisch

## Abstract

Recently, we demonstrated the development of a human pluripotent stem cell (hPSC)-derived adipocyte (fat cell) model with physiological insulin signaling. For the first time, a human in vitro adipocyte cell model has an insulin response similar to in vivo human adipocytes. We next leveraged this protocol to study the early stages of insulin resistance preceding Type 2 Diabetes, by exposing these cells to diabetic patient levels of hyperinsulinemia. This resulted in mild insulin resistance on all of the aforementioned phenotypes, reminiscent of early T2D development in humans. This is a powerful model to study the mechanisms of T2D development and the underlying molecular pathology. However, while adipocytes are an important metabolic cell type, insulin resistance and T2D are a whole-body affliction, involving many other cell types. To broaden the impact of our work and increase the excitement of the field for the study of insulin signaling and resistance, we expanded our protocol to a panel of 8 other hPSC-derived cell types. We demonstrate physiologically relevant insulin response for each cell type, and we highlight that our insulin-resistance inducing conditions blunt the insulin response in every cell type. Additionally, we validate that key cell identity markers are not lost during this adaptation. By adapting a simple, defined protocol to a wide panel of human in vitro cell types, we can now investigate the cell-autonomous origins of insulin resistance under physiologically relevant conditions.

## Main

Insulin resistance is one of the earliest signs of a panel of metabolic syndromes that includes Type 2 Diabetes (T2D), thereby serving as starting point for mechanistic study and opportunity for intervention. Insulin resistance is a disruption of normal insulin-mediated signaling, which typically manifests as connected metabolic disease phenotypes such as obesity, hyperinsulinemia, hyperglycemia, and hyperlipidemia. A mixture of diet, lifestyle, and genetics is known to contribute to the development of insulin resistance and its progression into T2D over time. Despite the public health significance of metabolic diseases, with hundreds of millions of T2D patients globally, human cell-based models for studying the nongenetic factors driving insulin resistance remain limited. As the in vitro results do not capture the potency and dose-response of in vivo insulin signaling, this results in an incomplete understanding of the cellular mechanisms driving human insulin resistance progression.

In a recent study published in Science Advances, we showcased a platform for mechanistic study of insulin signaling and resistance in an in vitro model using human pluripotent stem cell (hPSC)-derived adipocytes^1^. We specifically focused on making this in vitro human model relevant to physiological in vivo biology by using a culture medium that recapitulates the in vivo environment in terms of glucose and insulin levels. We performed the insulin sensitization and resistance induction experiments in hPSC-derived white and brown adipocytes in human embryonic and induced pluripotent stem cell lines, as well as in 3T3-L1 murine adipocytes and primary human stromal vascular fraction differentiated adipocytes^1^. Exposing these cells to a low-glucose DMEM-based medium with 0.1nM insulin for 5 days increases their insulin sensitivity, resulting in physiological insulin dose-response in AKT2 phosphorylation, insulin-stimulated glucose uptake, GLUT4 translocation, and suppression of lipolysis. Meanwhile, 5 days of 3nM hyperinsulinemia exposure induces insulin resistance in these cells, blunting the insulin response in all aforementioned assays. Through immunofluorescence, lipid measurements, and transcriptomic analysis we validated that hPSC-adipocytes do not lose their cell identity during this process.

However, insulin resistance is not limited to adipocytes and arises in a range of tissues, leading to pathogenesis throughout the body^2^. The insulin signaling pathway regulates cellular metabolism across a variety of cell types, yet insulin also has cell-type specific effects, for example insulin-stimulated glucose uptake in skeletal muscle and adipocytes, compared to insulin-suppression of gluconeogenesis and secretion in the liver^3^. To create a more comprehensive platform for studying metabolic disease, we sought to adapt our protocol to a diverse panel of hPSC-derived cell types Fig. 1a). As a methodological refinement, we transitioned from a DMEM-based medium to Human Plasma-Like Medium (HPLM), which more closely mimics the nutrient composition of human plasma and has been shown to improve the maturation of hPSC-derived cells^4^. We first validated this change by confirming that HPLM recapitulated our previous results in hPSC-adipocytes, and confirmed that both insulin sensitization and insulin resistance induction were identical with regards to AKT2 phosphorylation in hPSC-adipocytes between low-glucose DMEM (Friesen et al., 2022) and HPLM (Fig. 1b). This inspired confidence that we would be able to adapt an HPLM-based insulin-sensitization and resistance-inducing protocol to a panel of other hPSC-based cell types, which will unlock the mechanistic study of insulin signaling and resistance in the relevant cell types that contribute to whole-body T2D pathology.

**Fig. 1.**
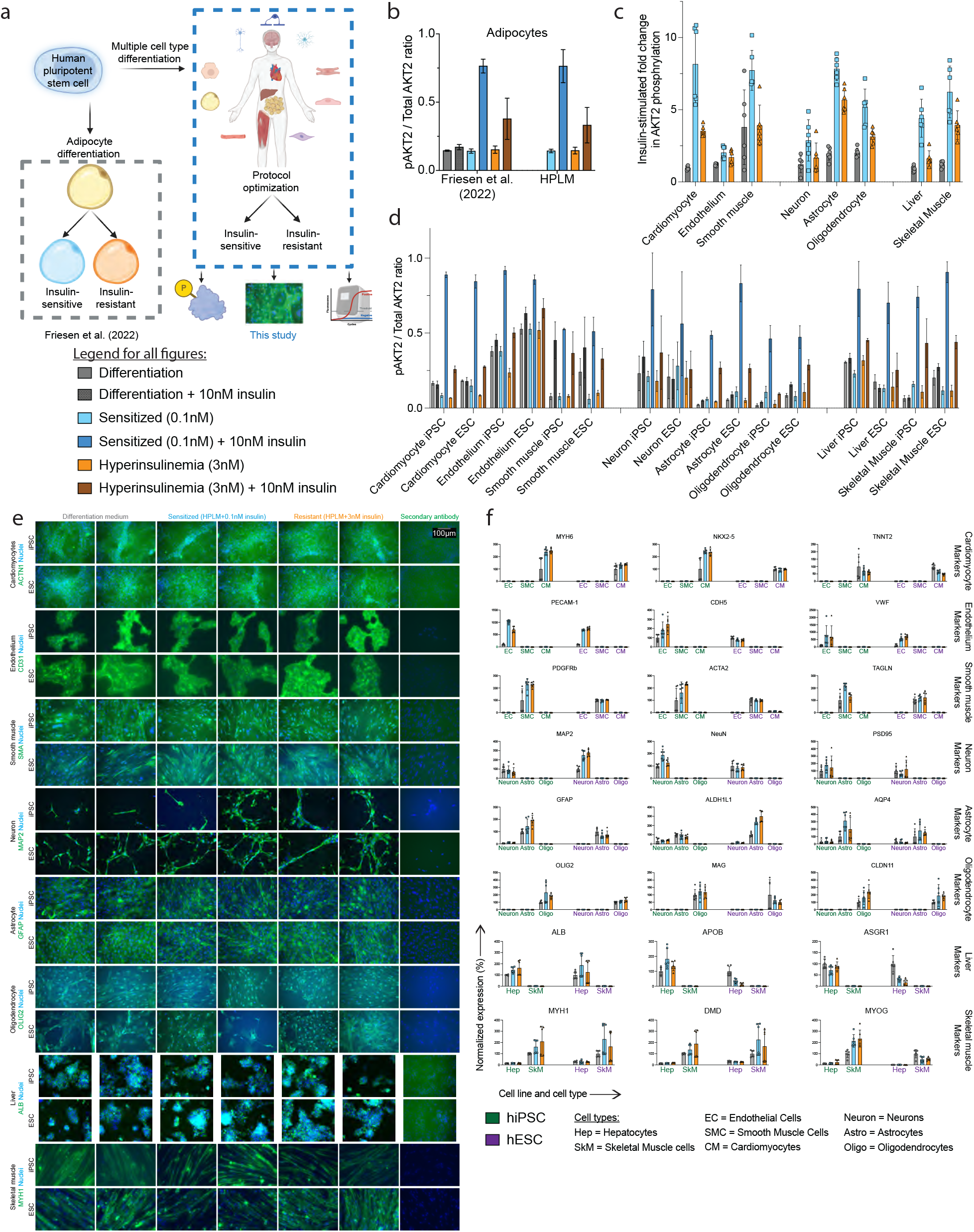
A broadly applicable protocol induces physiological insulin signaling and resistance across a panel of hPSC-derived cell types. a Schematic diagram illustrating the experimental design. Previously we generated insulin-sensitive and -resistant hPSC-adipocytes, in this study we adapted this protocol to a wide panel of hPSC-derived cell types. We test AKT2 protein phosphorylation and key cell identity marker expression through immunofluorescence and qPCR. b The results of the published protocol can be recapitulated by switching hPSC-adipocytes to HPLM. Shown is the ratio of phosphorylated over total AKT2 as quantified by ELISA (meanslll±lllSD; nlll=lll5 biological replicates, averaged from 2 technical ELISA replicates). c An adapted HPLM-based protocol leads to insulin sensitization compared to published differentiation medium, and blunted insulin response in hyperinsulinemia-induced insulin resistance. Shown is the acute insulin-stimulated fold change in AKT2 phosphorylation for each condition, normalized to total AKT2 protein as measured by ELISA (meanslll±lllSD, showing individual samples; combined results from nlll=lll3 replicate differentiations, averaged from 2 biological replicates, each averaged from 2 technical ELISA replicates, combined results from both an hiPSC and hESC line). d Data as in c, but split out between the hiPSC and hESC line, showing insulin sensitization and resistance across all cell types. Shown is the ratio of phosphorylated over total AKT2 as quantified by ELISA (meanslll±lllSD; nlll=lll3 replicate differentiations, averaged from 2 biological replicates, each averaged from 2 technical ELISA replicates). e Immunofluorescence imaging demonstrating unchanged key cell type marker expression of each cell type. Shown is one canonical marker for each cell type for each of the three conditions, and a secondary antibody control for both the hiPSC and hESC line (n = 2 replicate differentiations). f No deleterious impact on expression of cell type identity markers with our protocol as shown by qPCR. Shown are three canonical markers per cell type for each of the cell types, including other cell types in the respective groups (cardiovascular, brain, metabolic organ) as negative controls, for both the hiPSC and hESC line (meanslll±lllSD; nlll=lll3 replicate differentiations with 2 biological replicates each, averaged from 3 technical qPCR replicates each. Results are normalized to the housekeeping gene RPLP0 for each sample, after which each cell line is normalized to the differentiation condition).

We then differentiated a variety of hPSC-derived cell types using published protocols, including cells from the cardiovascular system (cardiomyocytes, endothelium, smooth muscle), the brain (neurons, astrocytes, oligodendrocytes), and metabolic organs (hepatocytes, skeletal muscle). Following their respective differentiations, cells were cultured for 5 days in HPLM with sensitizing (0.1nM) or resistance-inducing (3nM) concentrations of insulin. Remarkably, this single, unified protocol proved effective at tuning insulin response across a diverse panel of cells with minimal modifications (outlined in the methods). In all eight cell types, the sensitization protocol restored a robust, acute insulin-stimulated increase in pAKT2, which was largely absent in cells maintained in their standard, often insulin-rich, differentiation media, and a blunted response in the insulin resistance setting (Fig. 1c). One exception is the smooth muscle exhibiting insulin response in the differentiation medium, which was expected given that this differentiation medium does not contain any insulin. These results were highly reproducible across two genetically distinct hPSC lines—one embryonic (H1) and one induced pluripotent stem cell line—demonstrating the generalizability of the method (Fig. 1d). This showcases the potential of our protocol to generate physiologically insulin-sensitive or -resistant cells in a wide panel of hPSC-derived cell types.

A critical aspect of any disease modeling protocol is ensuring that the induced phenotype does not arise from a loss of cellular identity. We therefore confirmed that the 5-day protocol did not adversely affect the differentiated state of the cells. Immunofluorescence for canonical protein markers showed consistent expression across differentiation, sensitized, and resistant conditions for all cell types (Fig. 1e). Furthermore, quantitative PCR analysis of three lineage-specific mRNA markers for each cell type confirmed stable expression and the absence of off-target marker expression, validating that cell identity was maintained throughout the protocol (Fig. 1f). Altogether this confirms that our protocol does not negatively affect the cell identity of any of the tested cell types.

In summary, this study establishes a powerful new tool for metabolic research. By adapting a simple, defined protocol to a wide panel of human in vitro cell types, we can now investigate the cell-autonomous origins of insulin resistance under physiologically relevant conditions. Furthermore, the simplicity of our method ensures its broad adaptability to any hPSC-derived cell types, serving as a springboard for the field to create new models of metabolic disease. Critically, the use of a single base medium is a key advantage that paves the way for future studies on multi-tissue co-culture platforms designed to unravel the complex inter-organ communication that governs systemic metabolism.

## Data availability

All data and materials supporting this study are available from the corresponding author upon reasonable request. All antibodies used in this study are commercially available, and qPCR primers are listed in the methods.

## Acknowledgements

We thank A. Khalil, T. Lee, and J. Landtved for helpful comments. Figure 1a was created with the help of Biorender.com.

## Notes

### Competing Interest Statement

The authors have declared no competing interest.

## References

1 Friesen, M. et al. Development of a physiological insulin resistance model in human stem cell-derived adipocytes. Science Advances 8, eabn7298(2022). 10.1126/sciadv.abn7298

2 Rask-Madsen, C. & Kahn, C. R. Tissue–specific insulin signaling, metabolic syndrome, and cardiovascular disease. Arteriosclerosis, thrombosis, and vascular biology 32, 2052–2059 (2012).

3 James, D. E., Stöckli, J. & Birnbaum, M. J. The aetiology and molecular landscape of insulin resistance. Nature Reviews Molecular Cell Biology 22, 751–771 (2021). 10.1038/s41580-021-00390-6

4 Zhang, X. et al. Human Plasma-Like Medium Promotes Maturation of Human Pluripotent Stem Cell-Derived Cardiomyocytes. bioRxiv, 2025.2004. 2024.650456 (2025).

